# Genomic correlates for migratory direction in a free-ranging cervid

**DOI:** 10.1101/2022.05.26.493590

**Authors:** Maegwin Bonar, Spencer J. Anderson, Charles R. Anderson, George Wittemyer, Joseph M. Northrup, Aaron B. A. Shafer

## Abstract

Animal migrations are some of the most ubiquitous and one of the most threatened ecological processes. A wide range of migratory behaviours occur in nature, and this behaviour is not uniform between and within species, where even individuals in the same population can exhibit differentiation. While the environment largely drives migratory behaviour, it is necessary to understand the genetic mechanisms influencing migration to elucidate the adaptive potential of migratory species to cope with novel conditions and adapt to environmental change. In this study, we identified genes associated with migratory traits by undertaking pooled genome-wide scans on a natural population of a facultatively migrating ungulate. We identified genomic regions associated with variation in migratory direction in the study population, including FTM1, a gene linked to the formation of lipids, and DPPA3, a gene linked to epigenetic modifications of the maternal line. Such a genetic basis for migratory traits can have implications on heritability of behaviour and the flexibility of individuals to change their behaviour in the face of stochastic changes in their environment.

## Introduction

Migration, defined as the seasonal movement between home ranges (Dingle, 2014), is critical for the persistence of species in variable environments, influences nutrient deposition across ecosystems, and affects seed dispersal (Bauer & Hoye, 2014; Subalusky, Dutton, Rosi, & Post, 2017). Migratory behaviour has been identified across all major branches of the animal kingdom and is a complex phenomenon that involves the interaction between environmental and genetic cues, heritability, and selection (Griswold, Taylor, & Norris, 2010). Migration is one of the most threatened ecological processes (Harris, Thirgood, Hopcraft, Cromsigt, & Berger, 2009), and declines in migratory behaviour due to climate change and habitat loss have been observed, leading to the reduction in the variation in migratory behaviours exhibited across species, populations, and individuals (Berger, 2004; Bolger, Newmark, Morrison, & Doak, 2008; Festa-Bianchet, Ray, Boutin, Côté, & Gunn, 2011).

A wide range of migratory behaviours occur in nature, ranging from short altitudinal migrations in birds and bats (Boyle, 2017; Mcguire & Boyle, 2013) to the annual ∼1000km journey of wildebeest across East Africa (Hopcraft et al., 2014; Mijele et al., 2016). Variation in migratory behaviour exists within species (Lundberg et al., 2017), populations (Cavedon et al., 2019; Spitz, Hebblewhite, Stephenson, & German, 2018), and among individuals (Duijns et al., 2017a; Merkle et al., 2019), and variation in migration propensity, timing, and movement patterns can have critical implications for ecosystem connectivity (Bauer & Hoye, 2014). While variation in migratory behaviour is largely driven by the environment, which has been the focus of most research on migratory behaviour (Finger et al., 2016; Macdonald, Patterson, Hague, & Ian, 2011), a significant amount of residual variation can be explained by genetics (Franchini et al., 2017; Prince et al., 2017). Researchers have identified genes associated with migration direction and timing primarily in birds (Bazzi, Cecere, et al., 2016; Bazzi, Galimberti, et al., 2016; Saino et al., 2015) and fishes (Hess, Zendt, Matala, & Narum, 2016), but large mammals remain less studied (e.g., (Northrup, Shafer, Anderson Jr., & Coltman, 2014)). Understanding the genetic mechanisms driving migration can help elucidate the adaptive potential of migratory species to environmental change and help uncover the evolutionary origins and constraints of migration. Furthermore, as migratory behaviours continue to disappear across the globe, we could see a reduction additive genetic variance influencing migratory behaviours.

Ungulates (hooved mammals) show substantial phenotypic variation in migratory behaviour. For example, variation in migration propensity occurs within partially migrating populations (Hebblewhite & Merrill, 2009). Variation in migration timing occurs within species, populations, or individuals, similarly to migration distance, with some species of ungulates migrating thousands of kilometers (Subalusky et al., 2017) while others migrate less than 100 km (Jakes et al., 2018; Sawyer, Middleton, Hayes, Kauffman, & Monteith, 2016). Studies have primarily attributed migration in ungulates to variation in the abiotic and social environment (Abraham, Upham, Damian-serrano, & Jesmer, 2022; Leblond, St-Laurent, & Côté, 2016; van Beest, Vander Wal, Stronen, & Brook, 2013). Recently, Gervais et al. (2020) found evidence that movement (i.e., speed) and space-use behaviours are highly heritable in roe deer (*Capreolus capreolus*), which suggests genetic underpinnings for migratory behaviour in ungulates.

This study aims to identify genes associated with migratory traits by undertaking pooled genome-wide scans on a natural population of migrating mule deer (*Odocoileus hemionus*).

Specifically, we are using a study system consisting of four groups of facultatively migrating mule deer in the Piceance Basin of northwestern Colorado, USA, that all share the same winter range but migrate to two geographically distinct summer ranges (Figure 1, Appendix S1). Using genetic samples extracted from the blood of radio-tracked individuals, we assess diverent genomic regions correlated with different migratory behavior using whole genome sequencing and identified candidate genes associated with migration direction (e.g., Northrup et al., 2014).

**Figure 1.**
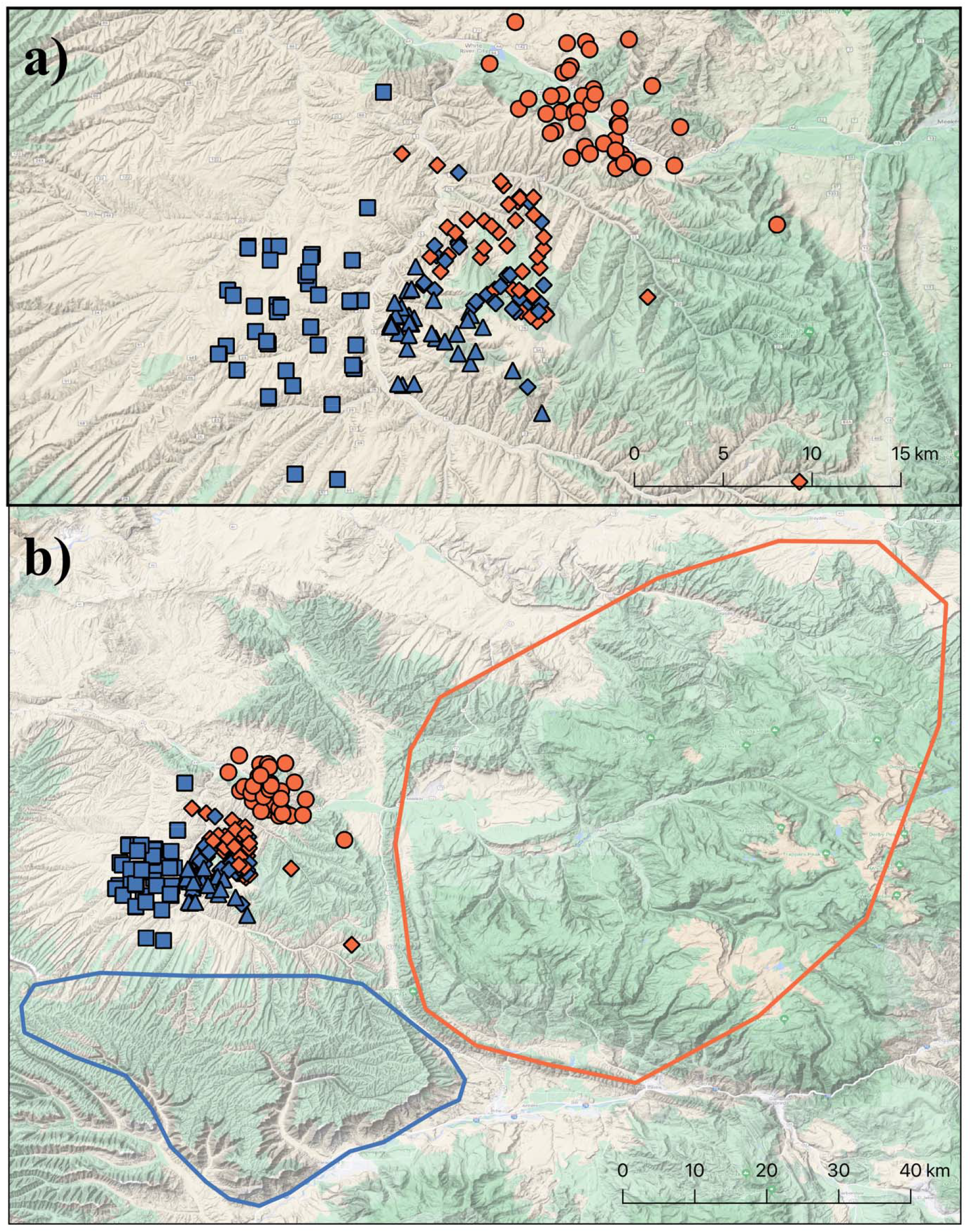
(a) winter range areas for Ryan Gulch (◼), South Magnolia (▲), North Magnolia (⍰), and North Ridge (•). Symbols represent study area, and colour represents migratory direction, with blue indicating north-south migration and orange indicating east-west migration. Points shown are the centroid of all GPS locations per individual while on the winter range. (b) Eastern and southern summer ranges are designated by the orange and blue outlines, respectively.

## Materials and Methods

### Data collection and migratory phenotypes

Mule deer were captured using a helicopter net gun on the four winter range study areas and fitted with store-on-board GPS radio collars (Advanced Telemetry Systems, Insanti MN, USA) with three different relocation schedules (5 hours, 60 minutes, and 30 minutes) depending on the individual. Deer were spotted visually by the helicopter capture crew and captured using a net gun. Deer were then blindfolded, hobbled, and administered 0.5 mg/kg of Midazolam (and 0.25 mg/kg of Azaperone intramuscularly) to alleviate capture-related stress (dose of both drugs based on an average weight of 75 kg). Deer were transported to a central processing site typically within 2 km of the capture site, where blood samples were collected for genetic analysis. Deer were released at the processing site immediately following blood sample collection and collar attachment. All procedures were approved by the Colorado State University (protocol ID: 10- 2350A) and Colorado Parks and Wildlife (protocol ID: 15-2008) Animal Care and Use Committees.

During spring migration, mule deer from the North Ridge and North Magnolia area move east-west towards the Flat Top Mountain Range, where they remain at high elevation for the summer. However, some mule deer from North Magnolia and the mule deer from the South Magnolia and Ryan Gulch populations migrate north-south to high elevations along the Roan Plateau (Lendrum, Jr, Monteith, Jenks, & Bowyer, 2013). Elevation on the winter study areas ranged from 1,675 to 2,285 m and from 2,000 to 2,800 m on the summer study areas. We visually examined the GPS data for each individual to determine which summer range each mule deer migrated to on the year they were collared. Mule deer tend to show high consistency and fidelity to their migratory routes (Northrup, Anderson, Gerber, & Wittemyer, 2021; Sawyer, Merkle, Middleton, Dwinnell, & Monteith, 2018). We then grouped individuals with the same migratory direction according to study group resulting in five groups as some individuals from North Magnolia migrate east-west while others migrate north-south (Figure 1). There is no genetic subdivision among populations (Northrup et al., 2014).

### DNA extraction and sequencing

We extracted DNA from each individual’s blood sample using the DNeasy Blood and Tissue Kit (Qiagen, Inc., Valencia, CA, USA), following the manufacturer’s protocol. We pooled the DNA of individuals from each of the five groups for sequencing using a pooled sequencing approach (Schlötterer, Tobler, Kofler, & Nolte, 2014). Equal quantities of DNA (100 ng/sample) were combined into representative pools (North Ridge (east-west migrating, n=50); North Magnolia (east-west migrating, n=50), North Magnolia (north-south migrating, n =33); South Magnolia (north-south migrating, n=50); and Ryan Gulch (north-south migrating, n=50) to a final concentration of 20 ng/ul of combined DNA for each pool. Each pool was sequenced across five lanes of an Illumina HiSeqX platform to a desired 50X coverage (total of 5 lanes) at The Centre for Applied Genomics (Toronto, ON) (Appendix S2, Table S1).

### Genome alignment and SNP calling

Mule deer and white-tailed deer can hybridize (Stubblefield, Warren, & Murphy, 1986), so we opted to use the long-read-based draft genome of white-tailed deer (*Odocoileus virginianus*) (Accession No. JAAVWD000000000) that was recently annotated (Anderson, Côté, Richard, & Shafer, 2022); we note that the available mule deer genome is simply a consensus sequence from reads mapped to earlier versions of the white-tailed reference (Russell et al., 2019). We performed initial quality filtering for all reads using fastqc and trimmed reads for quality and adaptors using the default Trimmomatic v.0.36 settings (Bolger et al. 2014). We aligned the pooled sequences to the unmasked white-tailed deer genome using BWA-mem v0.7.17 (Li, 2013). We used samtools v1.10 (Li et al., 2009) to merge and sort all aligned reads into five files representative of each pool. We filtered for duplicates using Picard v2.20.6 (Broad Institute, 2020), obtained uniquely mapped reads with samtools, and conducted local realignment using GATKv3.8 (Van der Auwera et al. 2013) before SNP calling. We called SNPs using samtools mpileup (parameters = -B -q20). We filtered out the masked regions and indels with 5bp flanking regions and removed all scaffolds <=50kb.

### Genome scan for population differentiation in migratory direction

We used an empirical F_ST_ approach (Akey et al., 2010; Cavedon et al., 2019) to identify potential SNPs relating to migratory phenotype between east-west-migrating and north-south-migrating pools. There is debate regarding the identification of outliers and the potential for false positives, specifically with regards to how the selection of an outlier cut-off level influences the process of identification (Lotterhos & Whitlock, 2014; Whitlock & Lotterhos, 2015). We used a sliding window approach in calculating F_ST_ using the Popoolation2 software suite (Kofler et al., 2011). We used 2500bp sliding widows with step sizes of 2500bp, and a minimum covered fraction of 0.8 and specified a minimum overall minor allele count of 3 for each pool. We had 5 pools that provided 10 pairwise comparisons, 6 opposite phenotype comparisons (east-west vs. north-south), and 4 same phenotype comparisons (1 east-west vs. east-west, 3 north-south vs. north-south). We defined an outlier window as being within the top 1% of F_ST_ values in at least 4/6 of the opposite phenotype comparisons and not within the top 1% of F_ST_ values in all 4 of the same phenotype comparisons. We also ran the same analysis using less conservative criteria and identified outlier windows from the top 1% of at least 3/6 of the opposite phenotype pairwise comparisons, and those results can be found in the supplementary material (Appendix S3).

We evaluated genetic differentiation between east-west-migrating and north-south-migrating pools by conducting a principal component analysis (PCA) using allele frequencies for the SNPs within outlier regions. We conducted one PCA using all available SNPs across the whole genome and one using the SNPs identified within the outlier regions identified through the F_ST_ approach. We expect that the PCA using the outlier SNPs putatively under selection, would show separation between the two migratory types, while the PCA using all SNPs would not. For all pools, we examined the proximity to genic regions for all SNPs identified within outlier regions using SnpEff v4.3t (Cingolani et al., 2012). For this, we generated VCF files using bcftools v1.9 (Li et al., 2009) and bed files containing the coordinates of all outlier SNPs. All SNP locations were characterized as being in an intergenic/intragenic region, 25 kb up/downstream of a gene (i.e., regulatory), intron, or exon. We also assessed the putative proximity to the nearest gene for every identified SNP.

### Gene ontology

To identify shared gene pathways among outlier SNPs, we used an analysis of gene ontology (GO) terms. We used the program Gowinda v1.12 (Kofler & Schlötterer, 2012) to determine GO term enrichment while accounting for gene length biases. We created a .gtf version of our annotation by removing duplicated genes, retaining only the longest version of each gene, which resulted in 15,395 unique genes in the annotation (Anderson et al., 2022). All outlier SNPs that SnpEff identified as being on or within 25kbp of genic regions were used to analyze gene ontology and compared to all outlier SNPs in every qualifying window. Finally, we plotted the GO results with WEGO (Web Gene Ontology Annotation Plot) to visualize annotations following the vocabularies and classifications provided by the GO Consortium (Ye et al., 2018).

## Results

### Detecting genetic variants and their genomic regions

After filtering for coverage, INDELS, and minimum allele counts, 194,533 windows each 2500bp wide were evaluated for each of the migratory-study area groups, respectively. The genome-wide mean F_ST_ value was 0.026, with 0.027 mean F_ST_ across each of the same migration direction comparisons and 0.026 mean F_ST_ across all opposite migratory group comparisons. Based on our pairwise comparisons, we identified 19 windows within the 99^th^ percentile in at least 4/6 pairwise comparisons of opposite migratory types and not within the 99^th^ percentile in the 4 same migratory type comparisons (Figure 2a). The average F_ST_ for these windows was 0.074 in the opposite migratory type comparisons and 0.027 in the same migratory type comparisons. Within these windows, 2903 SNPs were identified as on or within 25kbp of genic regions (Figure 2b). The PCA analysis using allele frequencies from the whole genome and outlier SNPs revealed separation of the migratory groups along the PC2 axis (Figure 2c).

**Figure 2.**
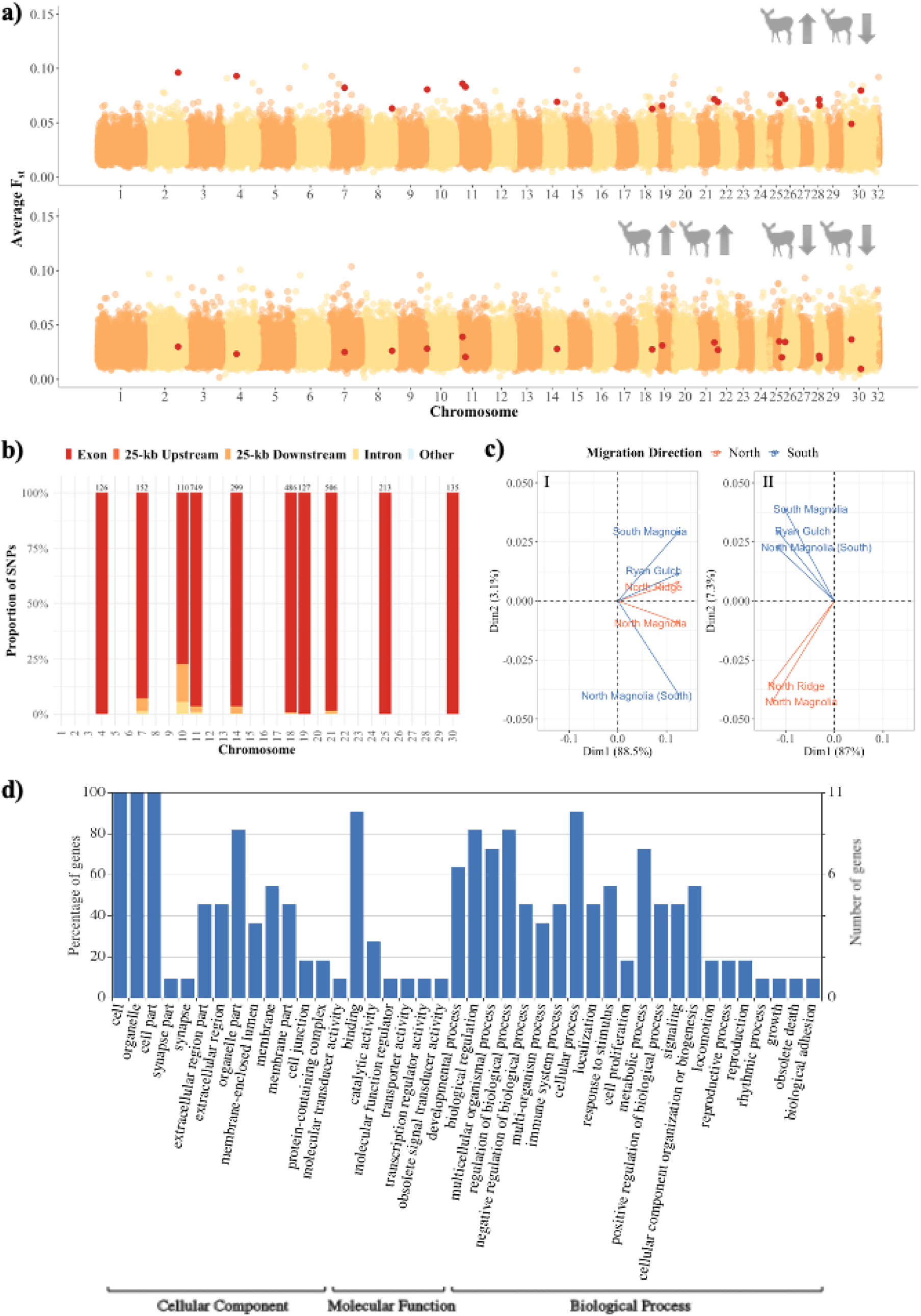
(a) Manhattan plot for average F_ST_ values across the whole genome, for all opposite migration comparisons (top) and same migration comparisons (bottom). Each point represents a 2500kbp window. The highlighted points indicate outlier windows associated with migratory direction, determined to have high levels of differentiation in at least 4/6 of the opposite migration comparisons and low levels of differentiation in all same migration comparisons. (b) Distribution of 2903 outlier SNPs across chromosomes identified as being on or within 25kbp of genic regions. Bars indicate the proportion of SNPs found on each chromosome distributed between exons, introns, 25kbp up- and downstream cites, and all other genic cites. Numbers above bars indicate the total number of outlier SNPs found on each chromosome. (c) Principal component analysis plot of major allele frequency values across 21,944,208 biallelic SNPs on the whole genome (I) and 463 biallelic outlier SNPs differentiated based on migratory direction (II). Arrows represent the 5 study group-migration direction groups with east-west migrators shown in orange and north-south migrators shown in blue. (d) Gene ontology (GO) assignment plots based on white-tailed deer annotations. Functional groups (x-axis) were in three functional categories: cellular component, molecular function, and biological process. Number and percent of genes within a given functional category performing a specific function are indicated on the y- axis. Some genes belong to more than one functional group, which may result in a sum exceeding 100% in a category. Plot was created using WEGO (Web Gene Ontology Annotation Plot; Ye et al., 2018).

### Gene Ontology Annotations

The results from GOWINDA on the non-intergenic outlier SNPs found on 11 genes, of note two of which included FTM1, a gene linked to the lipid synthesis in humans, and DPPA3, a gene linked to epigenetic modifications of the maternal line. GO analysis yielded 214 GO terms which were classified into 31 functional groups belonging to three functional categories: cellular components (11 groups), molecular functions (10 groups), and biological processes (10 groups) (Figure 2d, Table S2). Some genes belonged to more than one functional group (e.g., protein binding and cell differentiation), which sometimes resulted in a sum exceeding 100% in a category (e.g., cellular components). Among the genes categorized as cellular components, 100% were classified as cell parts. Most of the genes with molecular functions were associated with protein binding (90.9%), and most of the genes categorized as having biological processes were involved in cellular processes (90.9%), biological regulation (81.8%), and metabolic processes (72.7%).

## Discussion

Migratory behaviours continue to disappear globally, largely due to climate change and anthropogenic alteration to the environment (Wilcove & Wikelski, 2008). The disappearance of migratory routes can affect prey abundance, impact terrestrial and aquatic nutrient cycling, and could reduce genetic variation across species (Subalusky et al., 2017). When there is underlying additive genetic variance influencing migratory behaviours, a reduction in that variance can directly affect the adaptive potential of a species to cope with environmental change (Gervais et al., 2020). The heritability of migratory traits directly affects the response to selection and, given consistent directional selection on migration traits, could lead to distinct changes in migration variation within populations over a relatively short period of evolutionary time. Uncovering a genetic basis for migratory behaviour allows for genetic monitoring and could even be used as a measure to conserve biodiversity from individuals to whole ecosystems (Kardos, 2021). Our results found genomic regions associated with migration direction in mule deer; such a genetic basis for migratory traits can have implications for the heritability of this behaviour and the flexibility of individuals to change their behaviour in the face of stochastic changes to their environment.

The tiered window selection criteria seem to extract primarily functional outliers compared to other pooled genome-wide association studies using statistical thresholds (e.g., Anderson et al., 2020). The 11 genes that occurred on or within 25kb of a divergent window were ADGRB1, BSG, DPPA3, EMC9, FITM1, IKZF3, ITSN2, MAN2A2, PSME1, RNF216, and SETX. Of these 11 genes, ADGRB1, BSG, PSME1, SETX, RNF216, ITSN2, and IKZF3 are associated with immunity, host-virus interaction, and cell differentiation. The ability of an animal to maintain its immune system or mount an immune response can depend on its nutritional health and energetic condition. Variations in immune function and condition have been linked to migration in bats (Rogers et al., 2022) and stopover behaviour along a migration route in birds (Hegemann et al., 2018). MAN2A2, RNF216, and IKZF3 are involved in metal- and zinc-binding while EMC9 is part of the endoplasmic reticulum membrane protein complex. FITM1 is involved in lipid metabolism, and DPPA3 is involved in epigenetic chromatin reprogramming.

FTM1 is of note as it is related to the formation of lipid droplets and has been associated with fat storage in humans (Kadereit et al., 2008). Body fat is an indicator of fitness in ungulates (Bender, Lomas, & Browning, 2007). It can influence the annual survival of adult females (Tollefson, Shipley, Myers, Keisler, & Dasgupta, 2010), pregnancy and twinning rates (Johnstone-Yellin, Shipley, Myers, & Robinson, 2009), and the probability of a female rearing a fawn through summer (Cameron, Smith, Fancy, Gerhart, & White, 1993; Johnstone-Yellin et al., 2009). The link between body condition and migration performance has been documented in several taxa, including migratory shorebirds (Sjoerd Duijns et al., 2017), giant tortoises (Blake et al., 2013), and impalas (Gaidet & Lecomte, 2013)). The role of body fat in migration was suggested for this mule deer population previously (Northrup et al., 2014), supporting FTM1 as a potential modulator of mule deer migration.

A final noteworthy gene was DPPA3 that is related to epigenetic modification of the maternal germ cell line. Epigenetic variation can increase the phenotypic variation in offspring and thus increase the chance of offspring being able to cope with stochastic environments (Anastasiadi, Venney, Bernatchez, & Wellenreuther, 2021). Our dataset consisted entirely of female deer, and while ungulate migration behaviour is thought to have a learned component (e.g., white-tailed deer fawns follow their mother’s migration route; Nelson, 1998), previous work in this system has identified links between migratory timing and mitochondrial haplotype (Northrup et al., 2014). This, in conjunction with our results, could indicate that variation in migration behaviour is being inherited epigenetically through the maternal line. By validating these putative migration loci, it is conceivable that a gene panel could be developed for characterizing the genetic profiles of migrating populations and be used in wildlife management and recovery (Kardos, 2021).

It has been established that differences in migratory behaviour exist between genetically distinct populations (Cavedon et al., 2019; Sungani, Ngatunga, & Genner, 2016). Typically, genome-wide association studies of migratory behaviour using natural populations have two very distinct groups that are often geographically separated (Lundberg et al., 2017). We have shown that such large-scale comparisons are unnecessary to detect genomic variation associated with migratory phenotypes. We were able to identify distinct differences in the genomes of individuals who overlap in space on their winter range while migrating to two different summer ranges, demonstrating that genomic differentiation between migratory strategies is detectable at a fine scale. It is likely that similar detectable patterns may exist in other populations and species with intraindividual variation in migration behaviour. Many migratory ungulate species are considered at risk (e.g., caribou [*Rangifer tarandus*]), raising concerns regarding population isolation and loss of genetic diversity. Screening for similar genetic associations in more imperiled ungulate populations may help shed light on local population dynamics, heritability and evolvability, and better inform management decisions as migration routes continue to be affected by environmental and anthropogenic change.

## Acknowledgements

We respectfully acknowledge the territory where data were collected as the ancestral homelands of the Ute peoples. Data analyses were conducted at Trent University, which is on the traditional territory of the Mississauga Anishinaabeg.

This work was supported by Natural Sciences and Engineering Research Council of Canada (NSERC) Discovery Grant (ABAS and JMN); NSERC Vanier PhD Fellowship (MB); Compute Canada Resources for Research Groups (ABAS). Mule deer capture and monitoring was funded by Colorado Parks and Wildlife (CPW), White River Bureau of Land Management, ExxonMobil Production/XTO Energy, WPX Energy, Shell Exploration and Production, EnCana Corp., Marathon Oil Corp., Federal Aid in Wildlife Resto- ration (W-185-R), the Colorado Mule Deer Foundation, the Colorado Mule Deer Association, Safari Club International, Colorado Oil and Gas Conservation Commission, and the Colorado State Severance Tax. We thank L. Wolfe, C. Bishop, D. Finley (CPW) and numerous field technicians for capture expertise and field assistance. We thank Quicksilver Air, Inc. for deer captures, and L. Gepfert (CPW) and Coulter Aviation, Inc. for fixed-wing aircraft support.

## Data Accessibility

Data will be made available upon publication. Raw sequence data FastQ files will be available on the Sequence Read Archive (Accession: XXXXXX). All bioinformatic and analytical code available on GitLab (https://gitlab.com/WiDGeT_TrentU). Raw data will be uploaded to Dryad (Dryad Accession no. XXXXXX)

## Author Contributions

MB, SJA, ABAS, JMP designed the study; JMP, CRA, GW coordinated data collection and sample curation; MB performed research and analyzed data; MB wrote the manuscript with input from SJA, ABAS, CRA, GW, JMP.

## Notes

### Competing Interest Statement

The authors have declared no competing interest.

