## Supplementary Material for "Genomic correlates for migratory direction in a free-ranging cervid"

**Supplementary materials**

***Appendix S1:*** *Additional information on study area*

We used high-resolution GPS data from 233 adult female mule deer from four areas in the Piceance Basin of northwestern Colorado: North Ridge (53 km^2^); North Magnolia (79 km^2^); South Magnolia (83 km^2^); and Ryan Gulch (141 km^2^) (Figure S1). Climate in the region is characterized by warm dry summers and cold winters (Western Regional Climate Centre). Vegetation on the winter range is dominated by Pinion pine (*Pinus edulis*) and Utah juniper (*Juniperus osteosperma*), with the summer range being characterized by quaking aspen (*Populus tremuloides*), Douglas-fir (*Pseudotsuga menziesii*) forest, and Engelmann spruce (*Picea engelmannii*) subalpine fir (*Abies lasiocarpa*) forest at higher elevations (Garrott, White, Bartmann, Carpenter, & Alldredge, 1987; Northrup, Shafer, Anderson Jr., & Coltman, 2014).

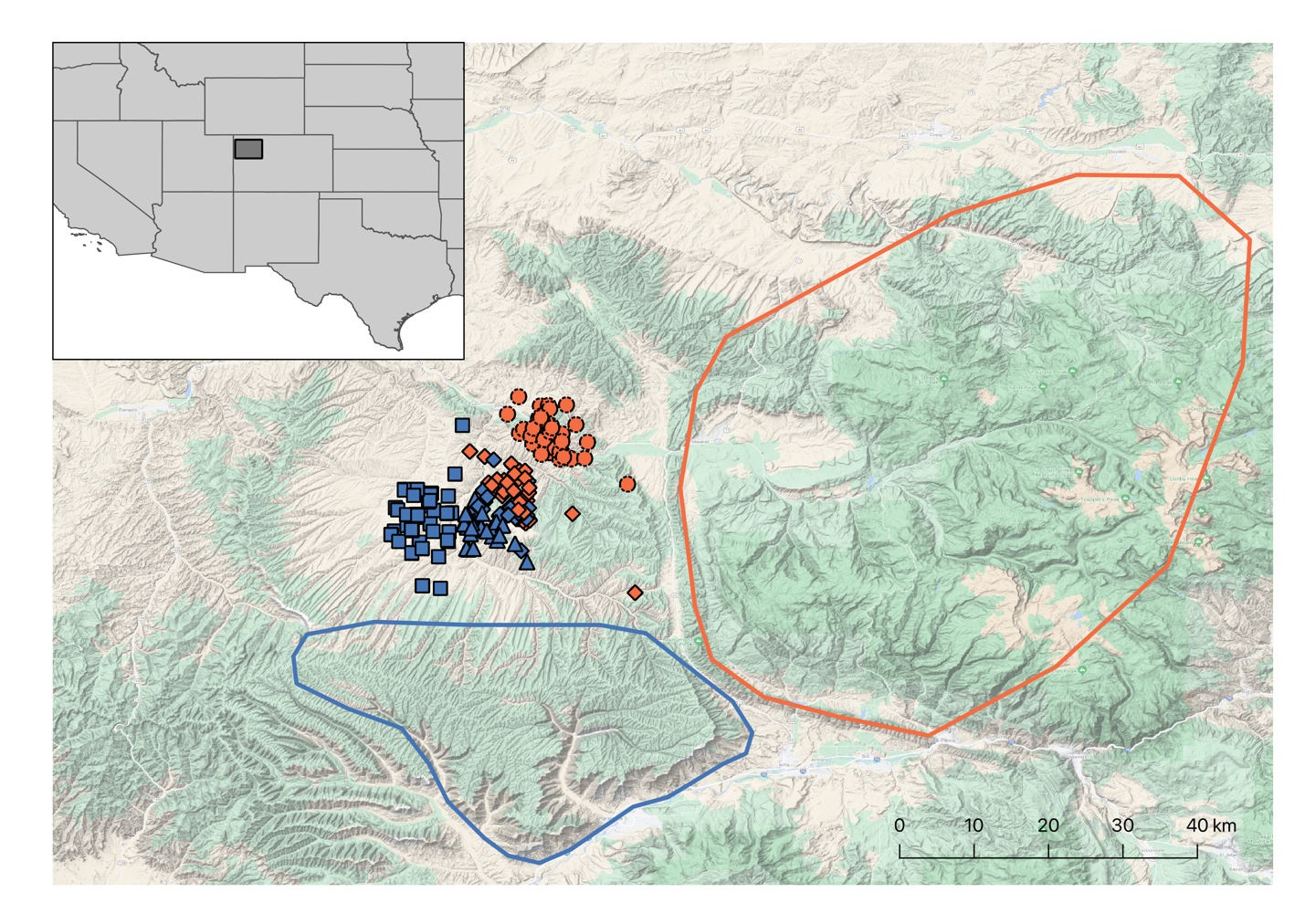

**Figure S1:** Study areas in the Piceance Basin of northwestern Colorado, USA: North Ridge (●, 53 km^2^); North Magnolia (◆, 79 km^2^); South Magnolia (▲, 83 km^2^); and Ryan Gulch (■, 141 km^2^). Symbols represent study group and colour represents migratory direction with blue indicating north-south migration and orange indicating east-west migration. Points shown are the centroid of all GPS locations per individual while on the winter range. Eastern and southern summer ranges are designated by the orange and blue outlines respectively.

***Appendix S2:*** *Sequencing information*

**Table S1:** Number of reads remaining after each filtering step and final genome wide coverage for each pool.

|  | **Initial** | **Deduplicateded** | **Unique** | **Coverage** |
| --- | --- | --- | --- | --- |
| North Magnolia | 885,404,886 | 790,433,632 | 648,399,900 | 35.6 |
| North Magnolia (South) | 685,338,615 | 770,955,285 | 631,223,528 | 34.6 |
| North Ridge | 890,861,042 | 633,029,250 | 649,381,675 | 35.6 |
| Ryan Gulch | 879,559,090 | 785,577,726 | 514,932,256 | 35.4 |
| South Magnolia | 848,004,029 | 752,455,051 | 616,581,653 | 33.8 |

***Appendix S3:*** *Genome scan for population differentiation in migratory direction using less conservative outlier criteria*

We repeated the analysis explained in the main text using the same empirical F_ST_ approach (Akey et al., 2010; Cavedon et al., 2019) to identify potential SNPs relating to migratory phenotype between north-migrating and south-migrating pools, but with a less conservative outlier selection criteria. We defined an outlier window as one that was within the top 1% of F_ST_ values in at least 3/6 of the opposite phenotype comparisons and was not within the top 1% of F_ST_ values in all 4 of the same phenotype comparisons (Cavedon et al., 2019).

The analysis was identical as outlined in the main text. For all outlier windows, we examined the proximity to genic regions for all SNPs identified within outlier regions using SnpEff v4.3t (Cingolani et al., 2012). We evaluated genetic differentiation between east-west-migrating and north-south-migrating pools by conducting a principal component analysis (PCA) using allele frequencies for the SNPs within outlier regions. And finally identify shared gene pathways among outlier SNPs we used an analysis of gene ontology (GO) terms. We used the program Gowinda v1.12 (Kofler & Schlötterer, 2012) to determine GO term enrichment, while also accounting for biases in gene length.

**Detecting genetic variants and their genomic regions**

Based on our pairwise comparisons we identified 113 windows that were within the 99^th^ percentile in at least 3/6 pairwise comparisons of opposite migratory types and not within the 99^th^ percentile in the 4 same migratory type comparisons (Figure S2). The average F_ST_ for these windows was 0.063 in the opposite migratory type comparisons and 0.027 in the same migratory type comparisons.

Within the 113 outlier windows, 13,416 SNPs were identified as on or within 25kbp of genic regions. 551 on Exons, 3495 introns, 2294 25kbp downstream, 7076 25kbp upstream (Figure S3). These 13,416 SNPs were found on or near 26 genes. The PCA analysis using allele frequencies from the whole genome and outlier SNPs revealed separation of the migratory groups along the PC2 axis (Figure S4).

**Gene Ontology Annotations**

The results from GOWINDA on the 13,416 non-intergenic SNPs found on 26 genes, yielded 267 GO terms which were classified into 76 functional groups belonging to three functional categories: cellular component (26 groups), molecular function (26 groups), and biological process (24 groups) (Figure S5, Table S3). Some genes belonged to more than one functional group (e.g., protein binding and cell differentiation), which sometimes resulted in a sum exceeding 100% in a category (e.g., cellular component). Among the genes categorized as cellular components, 100 % were classified as cell parts. Most of the genes with molecular functions were associated with protein binding (92.3 %), and most of the genes categorized as having biological processes were involved in cellular processes (88.5 %), biological regulation (80.8%) and metabolic processes (76.9 %).

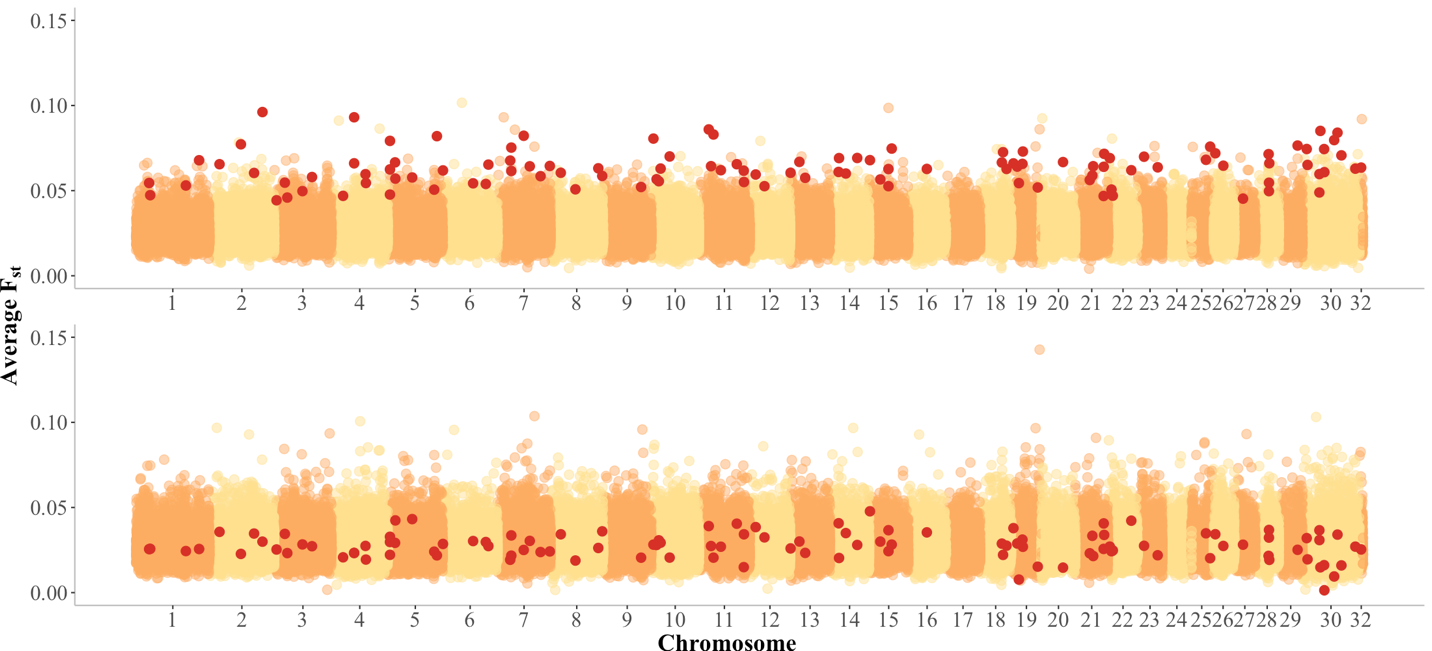

**Figure S2:** Manhattan plot for average F_ST_ values across the whole genome, for all opposite migration comparisons (top) and same migration comparisons (bottom). Each point represents a 2500kbp window, and points highlighted in red indicate outlier windows associated with migratory direction, determined to have high levels of differentiation in at least 3/6 of the opposite migration comparisons and low levels of differentiation in all same migration comparisons.

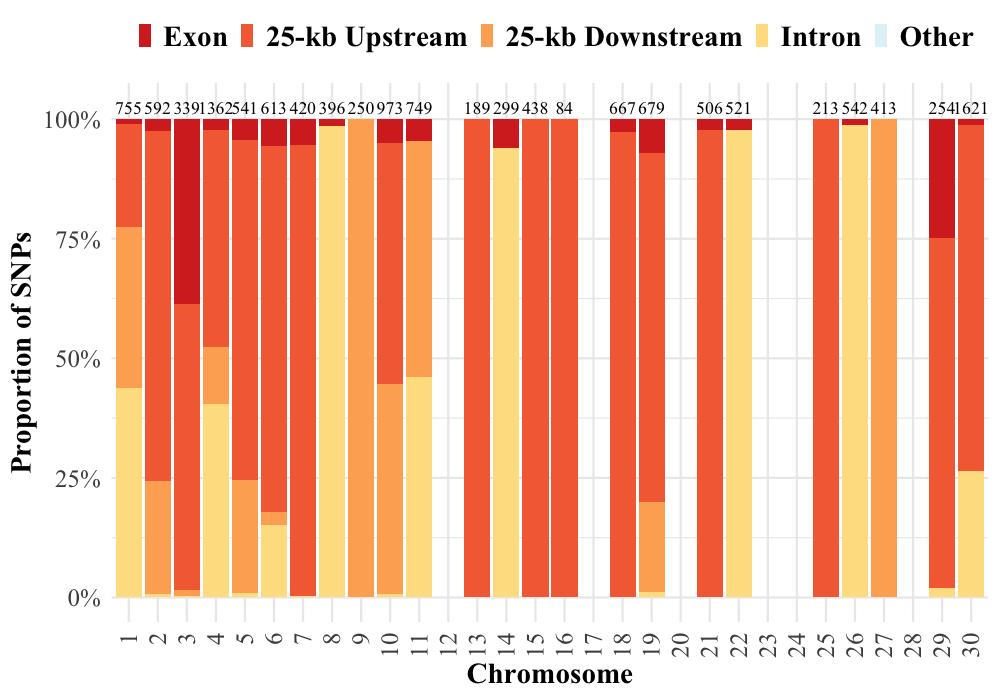

**Figure S3:** Distribution of 13,416 outlier SNPs across chromosomes identified as being on or within 25kbp of genic regions. Bars indicate the proportion of SNPs found on each chromosome distributed between exons, introns, 25kbp up- and downstream cites, and all other genic cites. Numbers above bars indicate the total number of outlier SNPs found on each chromosome.

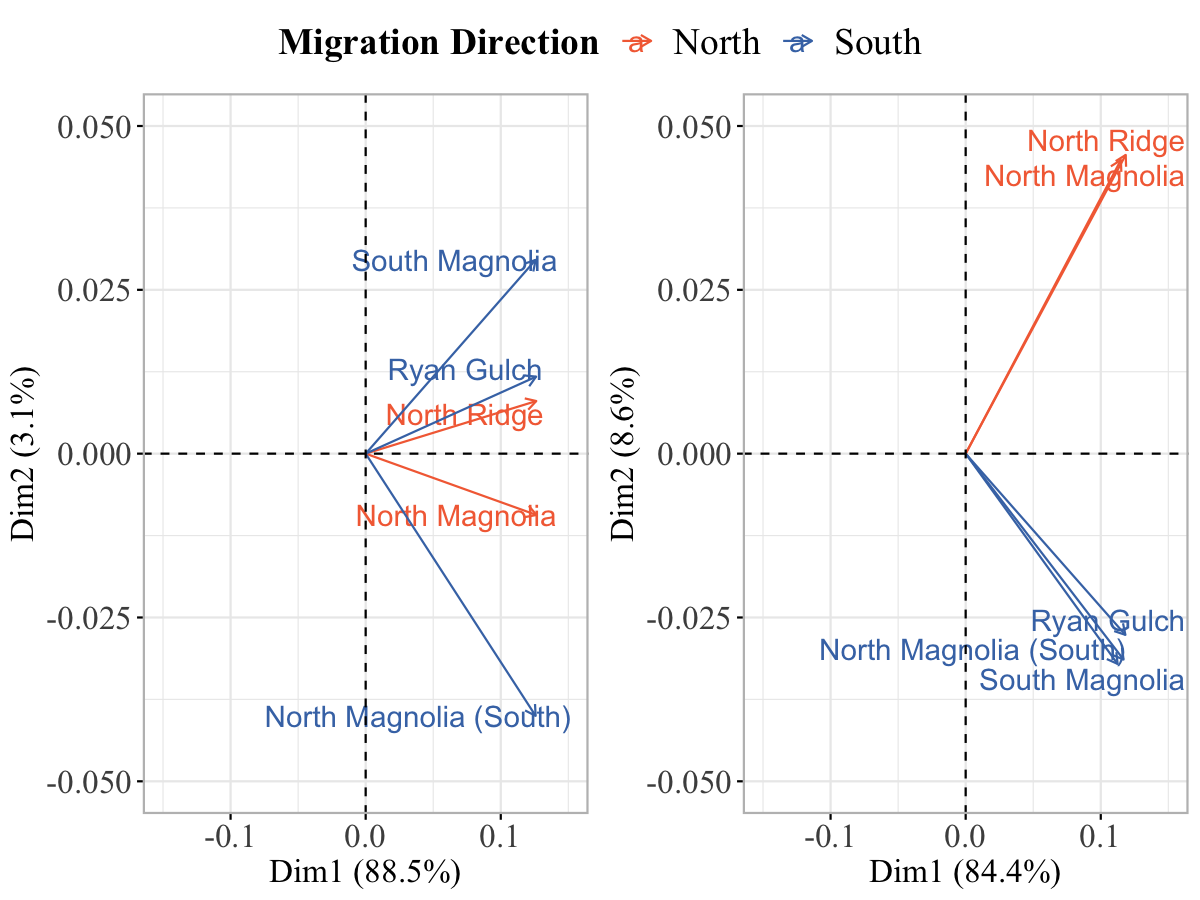

**Figure S4:** Principal component analysis plot of major allele frequency values across 21,944,208 biallelic SNPs on the whole genome (I) and 2935 biallelic outlier SNPs that are differentiated based on migratory direction (II). Arrows represent the 5 study group-migration direction groups with east-west migrators shown in orange and north-south migrators shown in blue.

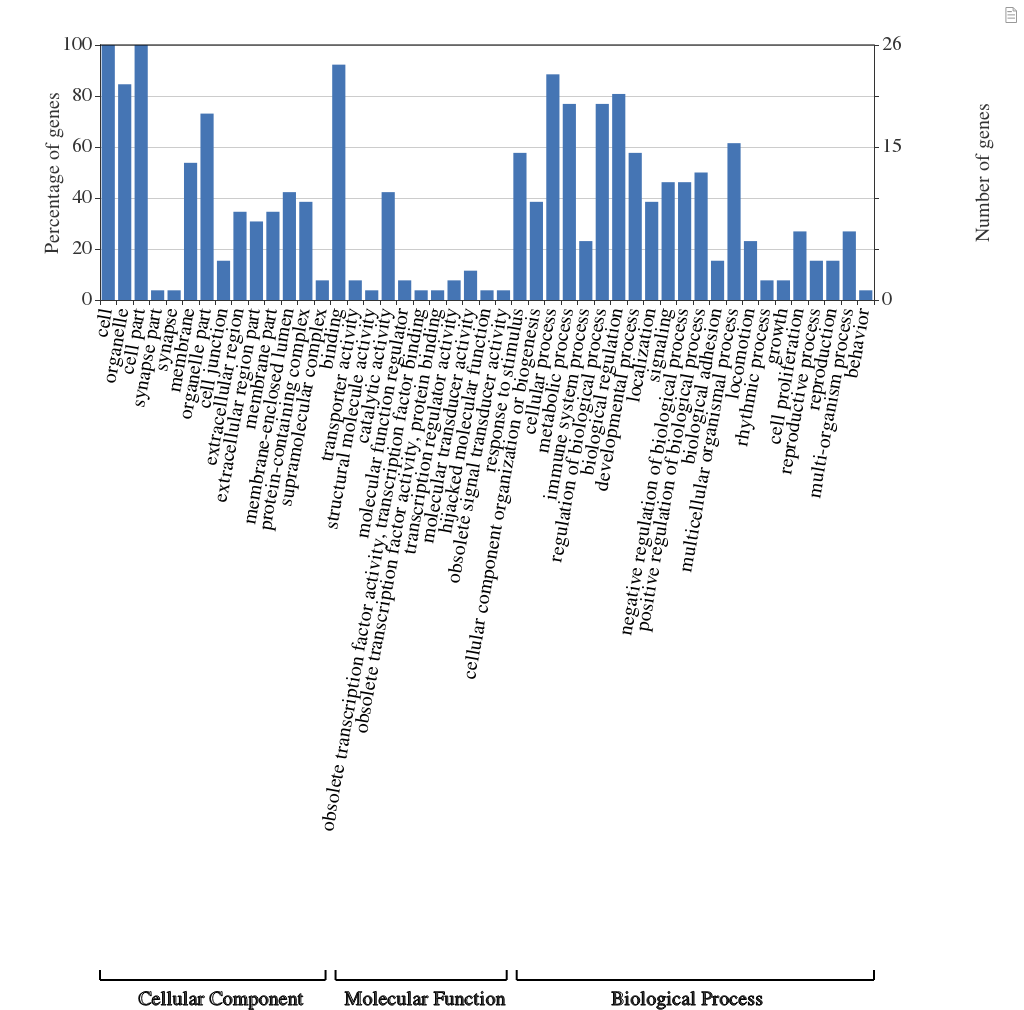

**Figure S5:** Gene ontology assignment plots based on white-tailed deer annotations. Functional groups (x-axis) were in three functional categories: cellular component, molecular function, and biological process. Number and percent of genes within a given functional category performing a specific function are indicated on the y-axis. Some genes belong to more than one functional group, which may result in a sum exceeding 100% in a category.

***Appendix S4:*** *Gene ontology analysis*

**Table S2:** Gene ontology (GO) tree showing the hierarchical breakdown of GO terms for SNPs within outlier windows found on genic regions. We defined an outlier window as one that was within the top 1% of F_ST_ values in at least 4/6 of the opposite phenotype comparisons and was not within the top 1% of F_ST_ values in all 4 of the same phenotype comparisons.

| **GO term** | **Gene number** | **Percentage** | **GO term description** | **Functional group** | **Functional category** |
| --- | --- | --- | --- | --- | --- |
| GO:0044464 | 11 | 100 | cell part | cell | Cellular component |
| GO:0099572 | 1 | 9.1 | postsynaptic specialization | organelle | Cellular component |
| GO:0043227 | 10 | 90.9 | membrane-bounded organelle | organelle | Cellular component |
| GO:0043230 | 3 | 27.3 | extracellular organelle | organelle | Cellular component |
| GO:0043229 | 10 | 90.9 | intracellular organelle | organelle | Cellular component |
| GO:0044422 | 9 | 81.8 | organelle part | organelle | Cellular component |
| GO:0043228 | 2 | 18.2 | non-membrane-bounded organelle | organelle | Cellular component |
| GO:0099572 | 1 | 9.1 | postsynaptic specialization | cell part | Cellular component |
| GO:0098794 | 1 | 9.1 | postsynapse | cell part | Cellular component |
| GO:0097458 | 2 | 18.2 | neuron part | cell part | Cellular component |
| GO:0044424 | 11 | 100 | intracellular part | cell part | Cellular component |
| GO:0005622 | 11 | 100 | intracellular | cell part | Cellular component |
| GO:0097223 | 1 | 9.1 | sperm part | cell part | Cellular component |
| GO:0012505 | 4 | 36.4 | endomembrane system | cell part | Cellular component |
| GO:0042175 | 2 | 18.2 | nuclear outer membrane-endoplasmic reticulum membrane network | cell part | Cellular component |
| GO:0042995 | 2 | 18.2 | cell projection | cell part | Cellular component |
| GO:0030427 | 1 | 9.1 | site of polarized growth | cell part | Cellular component |
| GO:0044463 | 2 | 18.2 | cell projection part | cell part | Cellular component |
| GO:0005886 | 3 | 27.3 | plasma membrane | cell part | Cellular component |
| GO:0071944 | 3 | 27.3 | cell periphery | cell part | Cellular component |
| GO:0044459 | 2 | 18.2 | plasma membrane part | cell part | Cellular component |
| GO:0098794 | 1 | 9.1 | postsynapse | synapse part | Cellular component |
| GO:0099572 | 1 | 9.1 | postsynaptic specialization | synapse part | Cellular component |
| GO:0097060 | 1 | 9.1 | synaptic membrane | synapse part | Cellular component |
| GO:0098984 | 1 | 9.1 | neuron to neuron synapse | synapse | Cellular component |
| GO:0044456 | 1 | 9.1 | synapse part | synapse | Cellular component |
| GO:0043230 | 3 | 27.3 | extracellular organelle | extracellular region part | Cellular component |
| GO:0045171 | 1 | 9.1 | intercellular bridge | extracellular region part | Cellular component |
| GO:0005615 | 4 | 36.4 | extracellular space | extracellular region part | Cellular component |
| GO:0044421 | 5 | 45.5 | extracellular region part | extracellular region | Cellular component |
| GO:0044446 | 9 | 81.8 | intracellular organelle part | organelle part | Cellular component |
| GO:0043233 | 4 | 36.4 | organelle lumen | organelle part | Cellular component |
| GO:0031090 | 3 | 27.3 | organelle membrane | organelle part | Cellular component |
| GO:0043233 | 4 | 36.4 | organelle lumen | membrane enclosed lumen | Cellular component |
| GO:0031090 | 3 | 27.3 | organelle membrane | membrane | Cellular component |
| GO:0098805 | 2 | 18.2 | whole membrane | membrane | Cellular component |
| GO:0042175 | 2 | 18.2 | nuclear outer membrane-endoplasmic reticulum membrane network | membrane | Cellular component |
| GO:0044425 | 5 | 45.5 | membrane part | membrane | Cellular component |
| GO:0005789 | 2 | 18.2 | endoplasmic reticulum membrane | membrane | Cellular component |
| GO:0098590 | 1 | 9.1 | plasma membrane region | membrane | Cellular component |
| GO:0005886 | 3 | 27.3 | plasma membrane | membrane | Cellular component |
| GO:0098589 | 2 | 18.2 | membrane region | membrane | Cellular component |
| GO:0005789 | 2 | 18.2 | endoplasmic reticulum membrane | membrane part | Cellular component |
| GO:0044459 | 2 | 18.2 | plasma membrane part | membrane part | Cellular component |
| GO:0098796 | 1 | 9.1 | membrane protein complex | membrane part | Cellular component |
| GO:0031224 | 4 | 36.4 | intrinsic component of membrane | membrane part | Cellular component |
| GO:0098589 | 2 | 18.2 | membrane region | membrane part | Cellular component |
| GO:0070161 | 2 | 18.2 | anchoring junction | cell junction | Cellular component |
| GO:0005911 | 1 | 9.1 | cell-cell junction | cell junction | Cellular component |
| GO:0030055 | 2 | 18.2 | cell-substrate junction | cell junction | Cellular component |
| GO:0098796 | 1 | 9.1 | membrane protein complex | protein-containing complex | Cellular component |
| GO:1902494 | 1 | 9.1 | catalytic complex | protein-containing complex | Cellular component |
| GO:0008537 | 1 | 9.1 | proteasome activator complex | protein-containing complex | Cellular component |
| GO:0022624 | 1 | 9.1 | proteasome accessory complex | protein-containing complex | Cellular component |
| GO:0038023 | 1 | 9.1 | signaling receptor activity | molecular transducer activity | Molecular function |
| GO:0060090 | 1 | 9.1 | molecular adaptor activity | binding | Molecular function |
| GO:0005515 | 9 | 81.8 | protein binding | binding | Molecular function |
| GO:1901363 | 2 | 18.2 | heterocyclic compound binding | binding | Molecular function |
| GO:0097159 | 2 | 18.2 | organic cyclic compound binding | binding | Molecular function |
| GO:0036094 | 3 | 27.3 | small molecule binding | binding | Molecular function |
| GO:0072341 | 1 | 9.1 | modified amino acid binding | binding | Molecular function |
| GO:0043167 | 6 | 54.5 | ion binding | binding | Molecular function |
| GO:0008289 | 1 | 9.1 | lipid binding | binding | Molecular function |
| GO:0097367 | 2 | 18.2 | carbohydrate derivative binding | binding | Molecular function |
| GO:0030246 | 2 | 18.2 | carbohydrate binding | binding | Molecular function |
| GO:0008144 | 1 | 9.1 | drug binding | binding | Molecular function |
| GO:0016787 | 2 | 18.2 | hydrolase activity | catalytic activity | Molecular function |
| GO:0140096 | 1 | 9.1 | catalytic activity, acting on a protein | catalytic activity | Molecular function |
| GO:0016740 | 1 | 9.1 | transferase activity | catalytic activity | Molecular function |
| GO:0140097 | 1 | 9.1 | catalytic activity, acting on DNA | catalytic activity | Molecular function |
| GO:0005085 | 1 | 9.1 | guanyl-nucleotide exchange factor activity | molecular function regulator | Molecular function |
| GO:0022857 | 1 | 9.1 | transmembrane transporter activity | transporter activity | Molecular function |
| GO:0003700 | 1 | 9.1 | DNA-binding transcription factor activity | transcription regulator activity | Molecular function |
| GO:0051093 | 1 | 9.1 | negative regulation of developmental process | developmental process | Biological process |
| GO:0048856 | 7 | 63.6 | anatomical structure development | developmental process | Biological process |
| GO:0050793 | 5 | 45.5 | regulation of developmental process | developmental process | Biological process |
| GO:0009653 | 4 | 36.4 | anatomical structure morphogenesis | developmental process | Biological process |
| GO:0048646 | 1 | 9.1 | anatomical structure formation involved in morphogenesis | developmental process | Biological process |
| GO:0051094 | 3 | 27.3 | positive regulation of developmental process | developmental process | Biological process |
| GO:0048869 | 5 | 45.5 | cellular developmental process | developmental process | Biological process |
| GO:0003006 | 1 | 9.1 | developmental process involved in reproduction | developmental process | Biological process |
| GO:0048589 | 1 | 9.1 | developmental growth | developmental process | Biological process |
| GO:0050789 | 9 | 81.8 | regulation of biological process | biological regulation | Biological process |
| GO:0065009 | 2 | 18.2 | regulation of molecular function | biological regulation | Biological process |
| GO:0065008 | 3 | 27.3 | regulation of biological quality | biological regulation | Biological process |
| GO:0007275 | 7 | 63.6 | multicellular organism development | multicellular organismal process | Biological process |
| GO:0051239 | 6 | 54.5 | regulation of multicellular organismal process | multicellular organismal process | Biological process |
| GO:0051241 | 2 | 18.2 | negative regulation of multicellular organismal process | multicellular organismal process | Biological process |
| GO:0051240 | 3 | 27.3 | positive regulation of multicellular organismal process | multicellular organismal process | Biological process |
| GO:0032504 | 2 | 18.2 | multicellular organism reproduction | multicellular organismal process | Biological process |
| GO:0001816 | 1 | 9.1 | cytokine production | multicellular organismal process | Biological process |
| GO:0044706 | 1 | 9.1 | multi-multicellular organism process | multicellular organismal process | Biological process |
| GO:0090130 | 1 | 9.1 | tissue migration | multicellular organismal process | Biological process |
| GO:0050793 | 5 | 45.5 | regulation of developmental process | regulation of biological process | Biological process |
| GO:0051239 | 6 | 54.5 | regulation of multicellular organismal process | regulation of biological process | Biological process |
| GO:0048519 | 5 | 45.5 | negative regulation of biological process | regulation of biological process | Biological process |
| GO:0043900 | 1 | 9.1 | regulation of multi-organism process | regulation of biological process | Biological process |
| GO:0050794 | 7 | 63.6 | regulation of cellular process | regulation of biological process | Biological process |
| GO:0002682 | 3 | 27.3 | regulation of immune system process | regulation of biological process | Biological process |
| GO:0019222 | 5 | 45.5 | regulation of metabolic process | regulation of biological process | Biological process |
| GO:0048518 | 5 | 45.5 | positive regulation of biological process | regulation of biological process | Biological process |
| GO:0048583 | 3 | 27.3 | regulation of response to stimulus | regulation of biological process | Biological process |
| GO:0023051 | 3 | 27.3 | regulation of signaling | regulation of biological process | Biological process |
| GO:0044087 | 2 | 18.2 | regulation of cellular component biogenesis | regulation of biological process | Biological process |
| GO:0032879 | 2 | 18.2 | regulation of localization | regulation of biological process | Biological process |
| GO:0040012 | 1 | 9.1 | regulation of locomotion | regulation of biological process | Biological process |
| GO:0040008 | 1 | 9.1 | regulation of growth | regulation of biological process | Biological process |
| GO:0051241 | 2 | 18.2 | negative regulation of multicellular organismal process | negative regulation of multicellular organismal process | Biological process |
| GO:0051093 | 1 | 9.1 | negative regulation of developmental process | negative regulation of multicellular organismal process | Biological process |
| GO:0023057 | 1 | 9.1 | negative regulation of signaling | negative regulation of multicellular organismal process | Biological process |
| GO:0040013 | 1 | 9.1 | negative regulation of locomotion | negative regulation of multicellular organismal process | Biological process |
| GO:0048523 | 4 | 36.4 | negative regulation of cellular process | negative regulation of multicellular organismal process | Biological process |
| GO:0048585 | 1 | 9.1 | negative regulation of response to stimulus | negative regulation of multicellular organismal process | Biological process |
| GO:0009892 | 3 | 27.3 | negative regulation of metabolic process | negative regulation of multicellular organismal process | Biological process |
| GO:0043900 | 1 | 9.1 | regulation of multi-organism process | multi-organism process | Biological process |
| GO:0051707 | 2 | 18.2 | response to other organism | multi-organism process | Biological process |
| GO:0044703 | 2 | 18.2 | multi-organism reproductive process | multi-organism process | Biological process |
| GO:0044419 | 1 | 9.1 | interspecies interaction between organisms | multi-organism process | Biological process |
| GO:0044706 | 1 | 9.1 | multi-multicellular organism process | multi-organism process | Biological process |
| GO:0044764 | 1 | 9.1 | multi-organism cellular process | multi-organism process | Biological process |
| GO:0019882 | 1 | 9.1 | antigen processing and presentation | immune system process | Biological process |
| GO:0006955 | 2 | 18.2 | immune response | immune system process | Biological process |
| GO:0002682 | 3 | 27.3 | regulation of immune system process | immune system process | Biological process |
| GO:0045321 | 1 | 9.1 | leukocyte activation | immune system process | Biological process |
| GO:0002684 | 1 | 9.1 | positive regulation of immune system process | immune system process | Biological process |
| GO:0002253 | 1 | 9.1 | activation of immune response | immune system process | Biological process |
| GO:0002252 | 1 | 9.1 | immune effector process | immune system process | Biological process |
| GO:0002520 | 2 | 18.2 | immune system development | immune system process | Biological process |
| GO:0050900 | 1 | 9.1 | leukocyte migration | immune system process | Biological process |
| GO:0050794 | 7 | 63.6 | regulation of cellular process | cellular process | Biological process |
| GO:0001775 | 1 | 9.1 | cell activation | cellular process | Biological process |
| GO:0044237 | 8 | 72.7 | cellular metabolic process | cellular process | Biological process |
| GO:0048522 | 4 | 36.4 | positive regulation of cellular process | cellular process | Biological process |
| GO:0051716 | 5 | 45.5 | cellular response to stimulus | cellular process | Biological process |
| GO:0006949 | 1 | 9.1 | syncytium formation | cellular process | Biological process |
| GO:0016043 | 6 | 54.5 | cellular component organization | cellular process | Biological process |
| GO:0140253 | 1 | 9.1 | cell-cell fusion | cellular process | Biological process |
| GO:0007154 | 5 | 45.5 | cell communication | cellular process | Biological process |
| GO:0007165 | 5 | 45.5 | signal transduction | cellular process | Biological process |
| GO:0048869 | 5 | 45.5 | cellular developmental process | cellular process | Biological process |
| GO:0006928 | 2 | 18.2 | movement of cell or subcellular component | cellular process | Biological process |
| GO:0048523 | 4 | 36.4 | negative regulation of cellular process | cellular process | Biological process |
| GO:0007049 | 1 | 9.1 | cell cycle | cellular process | Biological process |
| GO:0022402 | 1 | 9.1 | cell cycle process | cellular process | Biological process |
| GO:0016049 | 1 | 9.1 | cell growth | cellular process | Biological process |
| GO:0008219 | 3 | 27.3 | cell death | cellular process | Biological process |
| GO:0008037 | 1 | 9.1 | cell recognition | cellular process | Biological process |
| GO:0044764 | 1 | 9.1 | multi-organism cellular process | cellular process | Biological process |
| GO:0051641 | 1 | 9.1 | cellular localization | localization | Biological process |
| GO:0051234 | 4 | 36.4 | establishment of localization | localization | Biological process |
| GO:0032879 | 2 | 18.2 | regulation of localization | localization | Biological process |
| GO:0051674 | 2 | 18.2 | localization of cell | localization | Biological process |
| GO:0033036 | 2 | 18.2 | macromolecule localization | localization | Biological process |
| GO:0051235 | 1 | 9.1 | maintenance of location | localization | Biological process |
| GO:0006955 | 2 | 18.2 | immune response | response to stimulus | Biological process |
| GO:0009719 | 2 | 18.2 | response to endogenous stimulus | response to stimulus | Biological process |
| GO:0048583 | 3 | 27.3 | regulation of response to stimulus | response to stimulus | Biological process |
| GO:0048584 | 2 | 18.2 | positive regulation of response to stimulus | response to stimulus | Biological process |
| GO:0051716 | 5 | 45.5 | cellular response to stimulus | response to stimulus | Biological process |
| GO:0006950 | 4 | 36.4 | response to stress | response to stimulus | Biological process |
| GO:0042221 | 3 | 27.3 | response to chemical | response to stimulus | Biological process |
| GO:0009605 | 2 | 18.2 | response to external stimulus | response to stimulus | Biological process |
| GO:0009607 | 2 | 18.2 | response to biotic stimulus | response to stimulus | Biological process |
| GO:0009628 | 1 | 9.1 | response to abiotic stimulus | response to stimulus | Biological process |
| GO:0048585 | 1 | 9.1 | negative regulation of response to stimulus | response to stimulus | Biological process |
| GO:0070661 | 1 | 9.1 | leukocyte proliferation | cell proliferation | Biological process |
| GO:0042127 | 2 | 18.2 | regulation of cell proliferation | cell proliferation | Biological process |
| GO:0008285 | 1 | 9.1 | negative regulation of cell proliferation | cell proliferation | Biological process |
| GO:0009058 | 5 | 45.5 | biosynthetic process | metabolic process | Biological process |
| GO:0019222 | 5 | 45.5 | regulation of metabolic process | metabolic process | Biological process |
| GO:0044237 | 8 | 72.7 | cellular metabolic process | metabolic process | Biological process |
| GO:0071704 | 8 | 72.7 | organic substance metabolic process | metabolic process | Biological process |
| GO:0006807 | 7 | 63.6 | nitrogen compound metabolic process | metabolic process | Biological process |
| GO:0044238 | 7 | 63.6 | primary metabolic process | metabolic process | Biological process |
| GO:0009893 | 3 | 27.3 | positive regulation of metabolic process | metabolic process | Biological process |
| GO:0044281 | 2 | 18.2 | small molecule metabolic process | metabolic process | Biological process |
| GO:0009056 | 3 | 27.3 | catabolic process | metabolic process | Biological process |
| GO:0009892 | 3 | 27.3 | negative regulation of metabolic process | metabolic process | Biological process |
| GO:0032259 | 1 | 9.1 | methylation | metabolic process | Biological process |
| GO:0070085 | 1 | 9.1 | glycosylation | metabolic process | Biological process |
| GO:0070988 | 1 | 9.1 | demethylation | metabolic process | Biological process |
| GO:0009893 | 3 | 27.3 | positive regulation of metabolic process | positive regulation of biological process | Biological process |
| GO:0048522 | 4 | 36.4 | positive regulation of cellular process | positive regulation of biological process | Biological process |
| GO:0002684 | 1 | 9.1 | positive regulation of immune system process | positive regulation of biological process | Biological process |
| GO:0048584 | 2 | 18.2 | positive regulation of response to stimulus | positive regulation of biological process | Biological process |
| GO:0023056 | 2 | 18.2 | positive regulation of signaling | positive regulation of biological process | Biological process |
| GO:0051094 | 3 | 27.3 | positive regulation of developmental process | positive regulation of biological process | Biological process |
| GO:0044089 | 2 | 18.2 | positive regulation of cellular component biogenesis | positive regulation of biological process | Biological process |
| GO:0051240 | 3 | 27.3 | positive regulation of multicellular organismal process | positive regulation of biological process | Biological process |
| GO:0045927 | 1 | 9.1 | positive regulation of growth | positive regulation of biological process | Biological process |
| GO:1905954 | 1 | 9.1 | positive regulation of lipid localization | positive regulation of biological process | Biological process |
| GO:0023057 | 1 | 9.1 | negative regulation of signaling | signaling | Biological process |
| GO:0023051 | 3 | 27.3 | regulation of signaling | signaling | Biological process |
| GO:0023056 | 2 | 18.2 | positive regulation of signaling | signaling | Biological process |
| GO:0007165 | 5 | 45.5 | signal transduction | signaling | Biological process |
| GO:0007267 | 2 | 18.2 | cell-cell signaling | signaling | Biological process |
| GO:0016043 | 6 | 54.5 | cellular component organization | cellular component organization or biogenesis | Biological process |
| GO:0044085 | 2 | 18.2 | cellular component biogenesis | cellular component organization or biogenesis | Biological process |
| GO:0040012 | 1 | 9.1 | regulation of locomotion | locomotion | Biological process |
| GO:0048870 | 2 | 18.2 | cell motility | locomotion | Biological process |
| GO:0040013 | 1 | 9.1 | negative regulation of locomotion | locomotion | Biological process |
| GO:0003006 | 1 | 9.1 | developmental process involved in reproduction | reproductive process | Biological process |
| GO:0044703 | 2 | 18.2 | multi-organism reproductive process | reproductive process | Biological process |
| GO:0048609 | 2 | 18.2 | multicellular organismal reproductive process | reproductive process | Biological process |
| GO:0007566 | 1 | 9.1 | embryo implantation | reproductive process | Biological process |
| GO:0022414 | 2 | 18.2 | reproductive process | reproduction | Biological process |
| GO:0032504 | 2 | 18.2 | multicellular organism reproduction | reproduction | Biological process |
| GO:0019953 | 1 | 9.1 | sexual reproduction | reproduction | Biological process |
| GO:0007623 | 1 | 9.1 | circadian rhythm | rhythmic process | Biological process |
| GO:0040008 | 1 | 9.1 | regulation of growth | growth | Biological process |
| GO:0048589 | 1 | 9.1 | developmental growth | growth | Biological process |
| GO:0016049 | 1 | 9.1 | cell growth | growth | Biological process |
| GO:0045927 | 1 | 9.1 | positive regulation of growth | growth | Biological process |
| GO:0007155 | 1 | 9.1 | cell adhesion | biological adhesion | Biological process |

**Table S3:** Gene ontology (GO) tree showing the hierarchical breakdown of GO terms for SNPs within outlier windows found on genic regions. We defined an outlier window as one that was within the top 1% of F_ST_ values in at least 3/6 of the opposite phenotype comparisons and was not within the top 1% of F_ST_ values in all 4 of the same phenotype comparisons.

| **Go term** | **Gene number** | | **Percentage** | **GO term description** | **Functional group** | **Functional category** |
| --- | --- | --- | --- | --- | --- | --- |
| GO:0044464 | | 26 | 100 | cell part | cell | Cellular component |
| GO:0099572 | | 1 | 3.8 | postsynaptic specialization | organelle | Cellular component |
| GO:0043229 | | 21 | 80.8 | intracellular organelle | organelle | Cellular component |
| GO:0043227 | | 20 | 76.9 | membrane-bounded organelle | organelle | Cellular component |
| GO:0044422 | | 19 | 73.1 | organelle part | organelle | Cellular component |
| GO:0043228 | | 9 | 34.6 | non-membrane-bounded organelle | organelle | Cellular component |
| GO:0043230 | | 5 | 19.2 | extracellular organelle | organelle | Cellular component |
| GO:0099572 | | 1 | 3.8 | postsynaptic specialization | cell part | Cellular component |
| GO:0098794 | | 1 | 3.8 | postsynapse | cell part | Cellular component |
| GO:0097458 | | 3 | 11.5 | neuron part | cell part | Cellular component |
| GO:0044424 | | 24 | 92.3 | intracellular part | cell part | Cellular component |
| GO:0005622 | | 24 | 92.3 | intracellular | cell part | Cellular component |
| GO:0097223 | | 1 | 3.8 | sperm part | cell part | Cellular component |
| GO:0012505 | | 7 | 26.9 | endomembrane system | cell part | Cellular component |
| GO:0042995 | | 4 | 15.4 | cell projection | cell part | Cellular component |
| GO:0030427 | | 1 | 3.8 | site of polarized growth | cell part | Cellular component |
| GO:0044463 | | 3 | 11.5 | cell projection part | cell part | Cellular component |
| GO:0071944 | | 9 | 34.6 | cell periphery | cell part | Cellular component |
| GO:0005886 | | 8 | 30.8 | plasma membrane | cell part | Cellular component |
| GO:0031252 | | 1 | 3.8 | cell leading edge | cell part | Cellular component |
| GO:0044459 | | 5 | 19.2 | plasma membrane part | cell part | Cellular component |
| GO:0009986 | | 1 | 3.8 | cell surface | cell part | Cellular component |
| GO:1990204 | | 1 | 3.8 | oxidoreductase complex | cell part | Cellular component |
| GO:0042175 | | 1 | 3.8 | nuclear outer membrane-endoplasmic reticulum membrane network | cell part | Cellular component |
| GO:0043209 | | 1 | 3.8 | myelin sheath | cell part | Cellular component |
| GO:0044297 | | 1 | 3.8 | cell body | cell part | Cellular component |
| GO:0098794 | | 1 | 3.8 | postsynapse | synapse part | Cellular component |
| GO:0099572 | | 1 | 3.8 | postsynaptic specialization | synapse part | Cellular component |
| GO:0097060 | | 1 | 3.8 | synaptic membrane | synapse part | Cellular component |
| GO:0098984 | | 1 | 3.8 | neuron to neuron synapse | synapse | Cellular component |
| GO:0044456 | | 1 | 3.8 | synapse part | synapse | Cellular component |
| GO:0031090 | | 4 | 15.4 | organelle membrane | membrane | Cellular component |
| GO:0098805 | | 5 | 19.2 | whole membrane | membrane | Cellular component |
| GO:0005886 | | 8 | 30.8 | plasma membrane | membrane | Cellular component |
| GO:0044425 | | 9 | 34.6 | membrane part | membrane | Cellular component |
| GO:0098589 | | 4 | 15.4 | membrane region | membrane | Cellular component |
| GO:0098590 | | 1 | 3.8 | plasma membrane region | membrane | Cellular component |
| GO:0042175 | | 1 | 3.8 | nuclear outer membrane-endoplasmic reticulum membrane network | membrane | Cellular component |
| GO:0005789 | | 1 | 3.8 | endoplasmic reticulum membrane | membrane | Cellular component |
| GO:0044446 | | 19 | 73.1 | intracellular organelle part | organelle part | Cellular component |
| GO:0031090 | | 4 | 15.4 | organelle membrane | organelle part | Cellular component |
| GO:0043233 | | 11 | 42.3 | organelle lumen | organelle part | Cellular component |
| GO:0005911 | | 3 | 11.5 | cell-cell junction | cell junction | Cellular component |
| GO:0070161 | | 2 | 7.7 | anchoring junction | cell junction | Cellular component |
| GO:0030055 | | 2 | 7.7 | cell-substrate junction | cell junction | Cellular component |
| GO:0044421 | | 8 | 30.8 | extracellular region part | extracellular region | Cellular component |
| GO:0043230 | | 5 | 19.2 | extracellular organelle | extracellular region part | Cellular component |
| GO:0031012 | | 1 | 3.8 | extracellular matrix | extracellular region part | Cellular component |
| GO:0005615 | | 6 | 23.1 | extracellular space | extracellular region part | Cellular component |
| GO:0045171 | | 1 | 3.8 | intercellular bridge | extracellular region part | Cellular component |
| GO:0044420 | | 1 | 3.8 | extracellular matrix component | extracellular region part | Cellular component |
| GO:0098589 | | 4 | 15.4 | membrane region | membrane part | Cellular component |
| GO:0044459 | | 5 | 19.2 | plasma membrane part | membrane part | Cellular component |
| GO:0031224 | | 6 | 23.1 | intrinsic component of membrane | membrane part | Cellular component |
| GO:0098552 | | 1 | 3.8 | side of membrane | membrane part | Cellular component |
| GO:0098796 | | 2 | 7.7 | membrane protein complex | membrane part | Cellular component |
| GO:0005789 | | 1 | 3.8 | endoplasmic reticulum membrane | membrane part | Cellular component |
| GO:0043233 | | 11 | 42.3 | organelle lumen | membrane enclosed lumen | Cellular component |
| GO:1902494 | | 3 | 11.5 | catalytic complex | protein-containing complex | Cellular component |
| GO:0017053 | | 1 | 3.8 | transcriptional repressor complex | protein-containing complex | Cellular component |
| GO:0098796 | | 2 | 7.7 | membrane protein complex | protein-containing complex | Cellular component |
| GO:0036452 | | 1 | 3.8 | ESCRT complex | protein-containing complex | Cellular component |
| GO:0005667 | | 1 | 3.8 | transcription factor complex | protein-containing complex | Cellular component |
| GO:0016281 | | 1 | 3.8 | eukaryotic translation initiation factor 4F complex | protein-containing complex | Cellular component |
| GO:0008537 | | 1 | 3.8 | proteasome activator complex | protein-containing complex | Cellular component |
| GO:0022624 | | 1 | 3.8 | proteasome accessory complex | protein-containing complex | Cellular component |
| GO:0099081 | | 2 | 7.7 | supramolecular polymer | supramolecular complex | Cellular component |
| GO:0060090 | | 1 | 3.8 | molecular adaptor activity | binding | Molecular function |
| GO:0005515 | | 21 | 80.8 | protein binding | binding | Molecular function |
| GO:1901363 | | 7 | 26.9 | heterocyclic compound binding | binding | Molecular function |
| GO:0097159 | | 7 | 26.9 | organic cyclic compound binding | binding | Molecular function |
| GO:0072341 | | 1 | 3.8 | modified amino acid binding | binding | Molecular function |
| GO:0043167 | | 13 | 50 | ion binding | binding | Molecular function |
| GO:0048037 | | 2 | 7.7 | cofactor binding | binding | Molecular function |
| GO:0008144 | | 3 | 11.5 | drug binding | binding | Molecular function |
| GO:0036094 | | 6 | 23.1 | small molecule binding | binding | Molecular function |
| GO:0097367 | | 4 | 15.4 | carbohydrate derivative binding | binding | Molecular function |
| GO:0030246 | | 3 | 11.5 | carbohydrate binding | binding | Molecular function |
| GO:0044877 | | 2 | 7.7 | protein-containing complex binding | binding | Molecular function |
| GO:0008289 | | 1 | 3.8 | lipid binding | binding | Molecular function |
| GO:0042165 | | 1 | 3.8 | neurotransmitter binding | binding | Molecular function |
| GO:0022857 | | 2 | 7.7 | transmembrane transporter activity | transporter activity | Molecular function |
| GO:0140096 | | 6 | 23.1 | catalytic activity, acting on a protein | catalytic activity | Molecular function |
| GO:0016740 | | 2 | 7.7 | transferase activity | catalytic activity | Molecular function |
| GO:0016491 | | 2 | 7.7 | oxidoreductase activity | catalytic activity | Molecular function |
| GO:0016787 | | 8 | 30.8 | hydrolase activity | catalytic activity | Molecular function |
| GO:0140097 | | 1 | 3.8 | catalytic activity, acting on DNA | catalytic activity | Molecular function |
| GO:0016829 | | 1 | 3.8 | lyase activity | catalytic activity | Molecular function |
| GO:0005085 | | 2 | 7.7 | guanyl-nucleotide exchange factor activity | molecular function regulator | Molecular function |
| GO:0003700 | | 2 | 7.7 | DNA-binding transcription factor activity | transcription regulator activity | Molecular function |
| GO:0038023 | | 1 | 3.8 | signaling receptor activity | molecular transducer activity | Molecular function |
| GO:0001618 | | 1 | 3.8 | virus receptor activity | hijacked molecular function | Molecular function |
| GO:0042221 | | 8 | 30.8 | response to chemical | response to stimulus | Biological process |
| GO:0048583 | | 8 | 30.8 | regulation of response to stimulus | response to stimulus | Biological process |
| GO:0051716 | | 13 | 50 | cellular response to stimulus | response to stimulus | Biological process |
| GO:0009719 | | 3 | 11.5 | response to endogenous stimulus | response to stimulus | Biological process |
| GO:0048585 | | 2 | 7.7 | negative regulation of response to stimulus | response to stimulus | Biological process |
| GO:0006955 | | 4 | 15.4 | immune response | response to stimulus | Biological process |
| GO:0048584 | | 4 | 15.4 | positive regulation of response to stimulus | response to stimulus | Biological process |
| GO:0006950 | | 6 | 23.1 | response to stress | response to stimulus | Biological process |
| GO:0009628 | | 2 | 7.7 | response to abiotic stimulus | response to stimulus | Biological process |
| GO:0051606 | | 1 | 3.8 | detection of stimulus | response to stimulus | Biological process |
| GO:0009607 | | 2 | 7.7 | response to biotic stimulus | response to stimulus | Biological process |
| GO:0009605 | | 5 | 19.2 | response to external stimulus | response to stimulus | Biological process |
| GO:0016043 | | 10 | 38.5 | cellular component organization | cellular component organization or biogenesis | Biological process |
| GO:0044085 | | 5 | 19.2 | cellular component biogenesis | cellular component organization or biogenesis | Biological process |
| GO:0016043 | | 10 | 38.5 | cellular component organization | cellular process | Biological process |
| GO:0044237 | | 19 | 73.1 | cellular metabolic process | cellular process | Biological process |
| GO:0050794 | | 19 | 73.1 | regulation of cellular process | cellular process | Biological process |
| GO:0007154 | | 12 | 46.2 | cell communication | cellular process | Biological process |
| GO:0051716 | | 13 | 50 | cellular response to stimulus | cellular process | Biological process |
| GO:0007165 | | 11 | 42.3 | signal transduction | cellular process | Biological process |
| GO:0048523 | | 10 | 38.5 | negative regulation of cellular process | cellular process | Biological process |
| GO:0048522 | | 12 | 46.2 | positive regulation of cellular process | cellular process | Biological process |
| GO:0006949 | | 1 | 3.8 | syncytium formation | cellular process | Biological process |
| GO:0140253 | | 1 | 3.8 | cell-cell fusion | cellular process | Biological process |
| GO:0001775 | | 3 | 11.5 | cell activation | cellular process | Biological process |
| GO:0007049 | | 4 | 15.4 | cell cycle | cellular process | Biological process |
| GO:0022402 | | 3 | 11.5 | cell cycle process | cellular process | Biological process |
| GO:0008219 | | 4 | 15.4 | cell death | cellular process | Biological process |
| GO:0048869 | | 9 | 34.6 | cellular developmental process | cellular process | Biological process |
| GO:0006928 | | 5 | 19.2 | movement of cell or subcellular component | cellular process | Biological process |
| GO:0019725 | | 1 | 3.8 | cellular homeostasis | cellular process | Biological process |
| GO:0032940 | | 2 | 7.7 | secretion by cell | cellular process | Biological process |
| GO:0016049 | | 1 | 3.8 | cell growth | cellular process | Biological process |
| GO:0007163 | | 1 | 3.8 | establishment or maintenance of cell polarity | cellular process | Biological process |
| GO:0061919 | | 1 | 3.8 | process utilizing autophagic mechanism | cellular process | Biological process |
| GO:0030029 | | 1 | 3.8 | actin filament-based process | cellular process | Biological process |
| GO:0007059 | | 1 | 3.8 | chromosome segregation | cellular process | Biological process |
| GO:0051301 | | 1 | 3.8 | cell division | cellular process | Biological process |
| GO:0008037 | | 1 | 3.8 | cell recognition | cellular process | Biological process |
| GO:0044764 | | 1 | 3.8 | multi-organism cellular process | cellular process | Biological process |
| GO:0022412 | | 1 | 3.8 | cellular process involved in reproduction in multicellular organism | cellular process | Biological process |
| GO:0071704 | | 19 | 73.1 | organic substance metabolic process | metabolic process | Biological process |
| GO:0006807 | | 18 | 69.2 | nitrogen compound metabolic process | metabolic process | Biological process |
| GO:0044238 | | 18 | 69.2 | primary metabolic process | metabolic process | Biological process |
| GO:0044237 | | 19 | 73.1 | cellular metabolic process | metabolic process | Biological process |
| GO:0032259 | | 3 | 11.5 | methylation | metabolic process | Biological process |
| GO:0019222 | | 15 | 57.7 | regulation of metabolic process | metabolic process | Biological process |
| GO:0009058 | | 10 | 38.5 | biosynthetic process | metabolic process | Biological process |
| GO:0009892 | | 7 | 26.9 | negative regulation of metabolic process | metabolic process | Biological process |
| GO:0009893 | | 7 | 26.9 | positive regulation of metabolic process | metabolic process | Biological process |
| GO:0009056 | | 7 | 26.9 | catabolic process | metabolic process | Biological process |
| GO:0032963 | | 1 | 3.8 | collagen metabolic process | metabolic process | Biological process |
| GO:0044281 | | 4 | 15.4 | small molecule metabolic process | metabolic process | Biological process |
| GO:0055114 | | 2 | 7.7 | oxidation-reduction process | metabolic process | Biological process |
| GO:0070085 | | 1 | 3.8 | glycosylation | metabolic process | Biological process |
| GO:0070988 | | 1 | 3.8 | demethylation | metabolic process | Biological process |
| GO:0019882 | | 1 | 3.8 | antigen processing and presentation | immune system process | Biological process |
| GO:0002252 | | 2 | 7.7 | immune effector process | immune system process | Biological process |
| GO:0006955 | | 4 | 15.4 | immune response | immune system process | Biological process |
| GO:0002684 | | 2 | 7.7 | positive regulation of immune system process | immune system process | Biological process |
| GO:0002253 | | 2 | 7.7 | activation of immune response | immune system process | Biological process |
| GO:0002682 | | 3 | 11.5 | regulation of immune system process | immune system process | Biological process |
| GO:0045321 | | 3 | 11.5 | leukocyte activation | immune system process | Biological process |
| GO:0002520 | | 3 | 11.5 | immune system development | immune system process | Biological process |
| GO:0050900 | | 2 | 7.7 | leukocyte migration | immune system process | Biological process |
| GO:0031294 | | 1 | 3.8 | lymphocyte costimulation | immune system process | Biological process |
| GO:0050793 | | 9 | 34.6 | regulation of developmental process | regulation of biological process | Biological process |
| GO:0050794 | | 19 | 73.1 | regulation of cellular process | regulation of biological process | Biological process |
| GO:0023051 | | 9 | 34.6 | regulation of signaling | regulation of biological process | Biological process |
| GO:0048583 | | 8 | 30.8 | regulation of response to stimulus | regulation of biological process | Biological process |
| GO:0048519 | | 12 | 46.2 | negative regulation of biological process | regulation of biological process | Biological process |
| GO:0019222 | | 15 | 57.7 | regulation of metabolic process | regulation of biological process | Biological process |
| GO:0002682 | | 3 | 11.5 | regulation of immune system process | regulation of biological process | Biological process |
| GO:0048518 | | 13 | 50 | positive regulation of biological process | regulation of biological process | Biological process |
| GO:0044087 | | 3 | 11.5 | regulation of cellular component biogenesis | regulation of biological process | Biological process |
| GO:0051239 | | 9 | 34.6 | regulation of multicellular organismal process | regulation of biological process | Biological process |
| GO:0032879 | | 6 | 23.1 | regulation of localization | regulation of biological process | Biological process |
| GO:0040012 | | 3 | 11.5 | regulation of locomotion | regulation of biological process | Biological process |
| GO:0040008 | | 1 | 3.8 | regulation of growth | regulation of biological process | Biological process |
| GO:0030155 | | 1 | 3.8 | regulation of cell adhesion | regulation of biological process | Biological process |
| GO:0042752 | | 1 | 3.8 | regulation of circadian rhythm | regulation of biological process | Biological process |
| GO:0043900 | | 1 | 3.8 | regulation of multi-organism process | regulation of biological process | Biological process |
| GO:1900046 | | 1 | 3.8 | regulation of hemostasis | regulation of biological process | Biological process |
| GO:0050789 | | 20 | 76.9 | regulation of biological process | biological regulation | Biological process |
| GO:0065009 | | 7 | 26.9 | regulation of molecular function | biological regulation | Biological process |
| GO:0065008 | | 7 | 26.9 | regulation of biological quality | biological regulation | Biological process |
| GO:0050793 | | 9 | 34.6 | regulation of developmental process | developmental process | Biological process |
| GO:0009653 | | 8 | 30.8 | anatomical structure morphogenesis | developmental process | Biological process |
| GO:0048646 | | 4 | 15.4 | anatomical structure formation involved in morphogenesis | developmental process | Biological process |
| GO:0048856 | | 14 | 53.8 | anatomical structure development | developmental process | Biological process |
| GO:0051094 | | 6 | 23.1 | positive regulation of developmental process | developmental process | Biological process |
| GO:0048869 | | 9 | 34.6 | cellular developmental process | developmental process | Biological process |
| GO:0048589 | | 2 | 7.7 | developmental growth | developmental process | Biological process |
| GO:0051093 | | 3 | 11.5 | negative regulation of developmental process | developmental process | Biological process |
| GO:0003006 | | 2 | 7.7 | developmental process involved in reproduction | developmental process | Biological process |
| GO:0051641 | | 5 | 19.2 | cellular localization | localization | Biological process |
| GO:0051234 | | 8 | 30.8 | establishment of localization | localization | Biological process |
| GO:0033036 | | 3 | 11.5 | macromolecule localization | localization | Biological process |
| GO:0032879 | | 6 | 23.1 | regulation of localization | localization | Biological process |
| GO:0051674 | | 5 | 19.2 | localization of cell | localization | Biological process |
| GO:0051235 | | 1 | 3.8 | maintenance of location | localization | Biological process |
| GO:0007165 | | 11 | 42.3 | signal transduction | signaling | Biological process |
| GO:0023051 | | 9 | 34.6 | regulation of signaling | signaling | Biological process |
| GO:0023057 | | 2 | 7.7 | negative regulation of signaling | signaling | Biological process |
| GO:0023056 | | 4 | 15.4 | positive regulation of signaling | signaling | Biological process |
| GO:0007267 | | 3 | 11.5 | cell-cell signaling | signaling | Biological process |
| GO:0023057 | | 2 | 7.7 | negative regulation of signaling | negative regulation of biological process | Biological process |
| GO:0048585 | | 2 | 7.7 | negative regulation of response to stimulus | negative regulation of biological process | Biological process |
| GO:0048523 | | 10 | 38.5 | negative regulation of cellular process | negative regulation of biological process | Biological process |
| GO:0009892 | | 7 | 26.9 | negative regulation of metabolic process | negative regulation of biological process | Biological process |
| GO:0040013 | | 1 | 3.8 | negative regulation of locomotion | negative regulation of biological process | Biological process |
| GO:0051093 | | 3 | 11.5 | negative regulation of developmental process | negative regulation of biological process | Biological process |
| GO:0051241 | | 3 | 11.5 | negative regulation of multicellular organismal process | negative regulation of biological process | Biological process |
| GO:0034260 | | 1 | 3.8 | negative regulation of GTPase activity | negative regulation of biological process | Biological process |
| GO:0002684 | | 2 | 7.7 | positive regulation of immune system process | positive regulation of biological process | Biological process |
| GO:0048584 | | 4 | 15.4 | positive regulation of response to stimulus | positive regulation of biological process | Biological process |
| GO:0048522 | | 12 | 46.2 | positive regulation of cellular process | positive regulation of biological process | Biological process |
| GO:0023056 | | 4 | 15.4 | positive regulation of signaling | positive regulation of biological process | Biological process |
| GO:0051094 | | 6 | 23.1 | positive regulation of developmental process | positive regulation of biological process | Biological process |
| GO:0009893 | | 7 | 26.9 | positive regulation of metabolic process | positive regulation of biological process | Biological process |
| GO:0044089 | | 3 | 11.5 | positive regulation of cellular component biogenesis | positive regulation of biological process | Biological process |
| GO:0051240 | | 6 | 23.1 | positive regulation of multicellular organismal process | positive regulation of biological process | Biological process |
| GO:0045927 | | 1 | 3.8 | positive regulation of growth | positive regulation of biological process | Biological process |
| GO:0045785 | | 1 | 3.8 | positive regulation of cell adhesion | positive regulation of biological process | Biological process |
| GO:0051050 | | 2 | 7.7 | positive regulation of transport | positive regulation of biological process | Biological process |
| GO:1900048 | | 1 | 3.8 | positive regulation of hemostasis | positive regulation of biological process | Biological process |
| GO:0007155 | | 4 | 15.4 | cell adhesion | biological adhesion | Biological process |
| GO:0007275 | | 14 | 53.8 | multicellular organism development | multicellular organismal process | Biological process |
| GO:0051239 | | 9 | 34.6 | regulation of multicellular organismal process | multicellular organismal process | Biological process |
| GO:0032922 | | 1 | 3.8 | circadian regulation of gene expression | multicellular organismal process | Biological process |
| GO:0051240 | | 6 | 23.1 | positive regulation of multicellular organismal process | multicellular organismal process | Biological process |
| GO:0051241 | | 3 | 11.5 | negative regulation of multicellular organismal process | multicellular organismal process | Biological process |
| GO:0032504 | | 4 | 15.4 | multicellular organism reproduction | multicellular organismal process | Biological process |
| GO:0044706 | | 1 | 3.8 | multi-multicellular organism process | multicellular organismal process | Biological process |
| GO:0090130 | | 2 | 7.7 | tissue migration | multicellular organismal process | Biological process |
| GO:0035264 | | 1 | 3.8 | multicellular organism growth | multicellular organismal process | Biological process |
| GO:0022404 | | 1 | 3.8 | molting cycle process | multicellular organismal process | Biological process |
| GO:0042303 | | 1 | 3.8 | molting cycle | multicellular organismal process | Biological process |
| GO:0001816 | | 1 | 3.8 | cytokine production | multicellular organismal process | Biological process |
| GO:0050817 | | 2 | 7.7 | coagulation | multicellular organismal process | Biological process |
| GO:0040012 | | 3 | 11.5 | regulation of locomotion | locomotion | Biological process |
| GO:0048870 | | 5 | 19.2 | cell motility | locomotion | Biological process |
| GO:0040013 | | 1 | 3.8 | negative regulation of locomotion | locomotion | Biological process |
| GO:0052192 | | 1 | 3.8 | movement in environment of other organism involved in symbiotic interaction | locomotion | Biological process |
| GO:0042330 | | 1 | 3.8 | taxis | locomotion | Biological process |
| GO:0007623 | | 2 | 7.7 | circadian rhythm | rhythmic process | Biological process |
| GO:0007622 | | 1 | 3.8 | rhythmic behavior | rhythmic process | Biological process |
| GO:0040008 | | 1 | 3.8 | regulation of growth | growth | Biological process |
| GO:0048589 | | 2 | 7.7 | developmental growth | growth | Biological process |
| GO:0016049 | | 1 | 3.8 | cell growth | growth | Biological process |
| GO:0045927 | | 1 | 3.8 | positive regulation of growth | growth | Biological process |
| GO:0061351 | | 1 | 3.8 | neural precursor cell proliferation | cell proliferation | Biological process |
| GO:0042127 | | 7 | 26.9 | regulation of cell proliferation | cell proliferation | Biological process |
| GO:0008284 | | 3 | 11.5 | positive regulation of cell proliferation | cell proliferation | Biological process |
| GO:0070661 | | 1 | 3.8 | leukocyte proliferation | cell proliferation | Biological process |
| GO:0050673 | | 1 | 3.8 | epithelial cell proliferation | cell proliferation | Biological process |
| GO:0033687 | | 1 | 3.8 | osteoblast proliferation | cell proliferation | Biological process |
| GO:0008285 | | 3 | 11.5 | negative regulation of cell proliferation | cell proliferation | Biological process |
| GO:0044703 | | 4 | 15.4 | multi-organism reproductive process | reproductive process | Biological process |
| GO:0048609 | | 4 | 15.4 | multicellular organismal reproductive process | reproductive process | Biological process |
| GO:0009566 | | 1 | 3.8 | fertilization | reproductive process | Biological process |
| GO:0007566 | | 1 | 3.8 | embryo implantation | reproductive process | Biological process |
| GO:0003006 | | 2 | 7.7 | developmental process involved in reproduction | reproductive process | Biological process |
| GO:0022412 | | 1 | 3.8 | cellular process involved in reproduction in multicellular organism | reproductive process | Biological process |
| GO:0032504 | | 4 | 15.4 | multicellular organism reproduction | reproduction | Biological process |
| GO:0022414 | | 4 | 15.4 | reproductive process | reproduction | Biological process |
| GO:0019953 | | 3 | 11.5 | sexual reproduction | reproduction | Biological process |
| GO:0044703 | | 4 | 15.4 | multi-organism reproductive process | multiorganism process | Biological process |
| GO:0044419 | | 3 | 11.5 | interspecies interaction between organisms | multiorganism process | Biological process |
| GO:0051707 | | 2 | 7.7 | response to other organism | multiorganism process | Biological process |
| GO:0044706 | | 1 | 3.8 | multi-multicellular organism process | multiorganism process | Biological process |
| GO:0043900 | | 1 | 3.8 | regulation of multi-organism process | multiorganism process | Biological process |
| GO:0044764 | | 1 | 3.8 | multi-organism cellular process | multiorganism process | Biological process |
| GO:0007622 | | 1 | 3.8 | rhythmic behavior | behavior | Biological process |
| GO:0007626 | | 1 | 3.8 | locomotory behavior | behavior | Biological process |

**REFERERNCES**

Akey, J. M., Ruhe, A. L., Akey, D. T., Wong, A. K., Connelly, C. F., Madeoy, J., … Neff, M. W. (2010). Tracking footprints of artificial selection in the dog genome. *Proceedings of the National Academy of Sciences of the United States of America*, *107*(3), 1160–1165. doi: 10.1073/pnas.0909918107

Cavedon, M., Gubili, C., Heppenheimer, E., VonHoldt, B., Mariani, S., Hebblewhite, M., … Musiani, M. (2019). Genomics, environment and balancing selection in behaviourally bimodal populations: The caribou case. *Molecular Ecology*, *28*(8), 1946–1963. doi: 10.1111/mec.15039

Cingolani, P., Platts, A., Wang, L. L., Coon, M., Nguyen, T., Wang, L., … Ruden, D. M. (2012). A program for annotating and predicting the effects of single nucleotide polymorphisms, SnpEff: SNPs in the genome of Drosophila melanogaster strain w1118; iso-2; iso-3. *Fly*, *6*(2), 80–92. doi: 10.4161/fly.19695

Garrott, R. A., White, G. C., Bartmann, R. M., Carpenter, L. H., & Alldredge, A. W. (1987). Movements of Female Mule Deer in Northwest Colorado. *Journal of Wildlife Management*, *51*(3), 634–643.

Kofler, R., & Schlötterer, C. (2012). Gowinda: Unbiased analysis of gene set enrichment for genome-wide association studies. *Bioinformatics*, *28*(15), 2084–2085. doi: 10.1093/bioinformatics/bts315

Northrup, J. M., Shafer, A. B. A., Anderson Jr., C. R., & Coltman, D. W. (2014). Fine-scale genetic correlates to condition and migration in a wild cervid. *Evolutionary Applications*, *7*, 937–948. doi: 10.1111/eva.12189
